# EchoVisuALL: From Echocardiography to Gene Discovery

**DOI:** 10.64898/2026.02.18.706519

**Authors:** Isabella Galter, Elida Schneltzer, Carsten Marr, Nadine Spielmann, Martin Hrabě de Angelis

## Abstract

Cardiovascular diseases are a major global health burden, demanding phenotyping frameworks that can match the scale and complexity of contemporary mouse genetics. Here, we introduce EchoVisuALL, an AI-enabled pipeline for automated high-throughput transthoracic echocardiography (TTE) coupling deep-learning-based left-ventricular segmentation with data reporting. Across 65,000 recordings from over 18,000 mice, including single-gene knockouts from the International Mouse Phenotyping Consortium, the framework quantified cardiac morphology and function with minimal operator dependency and high reliability, validated against an expert-curated gold standard dataset. By extracting quantitative parameters across the cardiac cycle, EchoVisuALL in combination with multi-dimensional clustering uncovered nonlinear phenotypic relationships and revealed 37 of 715 genes associated with significant cardiac abnormalities, encompassing well-known human disease genes as well as 12 previously unrecognized candidates, including *Cep70, Acot12, Atp8b3, Eea1, Kctd2*, and *Tspan15*. These genotype-phenotype associations are involved in myocardial energetics, membrane biology, and cardiac remodeling. We demonstrate the potential of EchoVisuALL to move beyond image segmentation by delivering a standardized, quantitative foundation for scalable downstream analyses, enabling the discovery of novel cardiac disease genes.

## Introduction

Cardiovascular disease remains a major global health burden [1], yet its molecular basis is only partially understood [2]. Although genome-wide association studies (GWAS) have identified hundreds of loci, causal deduction remains challenging due to the complexity of cardiac genetics [3, 4, 5]. Experimental models with precise genetic manipulation, such as the mouse, whose four-chambered heart closely resembles human anatomy [6, 7] and whose developmental trajectories demonstrate a remarkable congruence to the human heart [8], are essential for dissecting mechanisms and causality in vivo [9, 10, 11, 12].

Transthoracic echocardiography (TTE) is the gold standard for non-invasive cardiac assessment in both clinical diagnostics and experimental research [13, 14]. Its ability to quantify cardiac structure and function makes TTE indispensable, but manual interpretation is labor-intensive and challenging at scale [15]. As datasets grow, especially in multi-center initiatives as the International Mouse Phenotyping Consortium (IMPC), manual analysis has become a major bottleneck, and subtle phenotypes risk remaining undetected [16].

AI-based image analysis offers a promising solution [17], enabling rapid and consistent extraction of cardiac metrics across a wide range of imaging modalities in patients [18, 19, 20, 21], with a particular emphasis on left-ventricular (LV) diagnostics [22], paralleling our investigation in mouse TTE. While several AI tools exist for human cardiac imaging [23], counterparts for murine imaging are far fewer and often lack generalizability because they are trained on narrowly constrained datasets [24, 25, 26, 27].

To address this limitation, we developed AI models tailored to the diversity of mouse TTE recordings. Echo2Pheno [16] introduced LV detection in awake TTE imaging, and EchoVisuAL [28] expanded segmentation performance across the full variability of the IMPC data. Yet segmentation alone cannot capture cardiac dysfunction; extracting measurements of cardiac parameters throughout the cardiac cycle is required to translate raw echocardiogram images into actionable data [29, 30].

EchoVisuALL addresses this limitation by integrating deep–learning–based LV segmentation with automated extraction of key structural and functional metrics. These encompass LV chamber volume and internal diameter as structural indices, and heart rate, ejection fraction, fractional shortening, stroke volume, and cardiac output as functional readouts. To demonstrate the capabilities of EchoVisuALL, we reanalyzed thousands of recordings from IMPC mice. By moving beyond univariate linear models and instead applying density-based multidimensional clustering together with wild-type control reference ranges, the framework identified gene-specific cardiac phenotypes in an unbiased, biologically grounded manner. Reframing large-scale TTE as a discovery platform rather than a manual evaluation task, EchoVisuALL uncovered genotype-phenotype relationships, including subtle previously unrecognized cardiac gene functions, and resolved a central bottleneck in comprehensive cardiac phenotyping.

## Results

Our results are based on a large-scale analysis comprising 65,241 transthoracic echocardiography (TTE) recordings from 18,402 mice. Using a newly developed segmentation model, EchoVisuALL, we achieved automated and standardized evaluation of all recordings. These functional readouts provided the foundation for a multidimensional, density-based clustering analysis. This large-scale analysis identified genes previously unrecognized in cardiac pathology while also confirming known associations with heart disease. To ensure robustness and interpretability, a curated gold-standard dataset was established and used to benchmark model performance and reliability.

### Benchmark dataset establishes a reliable gold standard

To validate EchoVisuALL’s automated evaluation, we established an expert-curated gold standard dataset consisting of 20 manually annotated TTE recordings with a total of 836 frames. Five independent experts provided manual LV segmentations, which were then aggregated into a consensus reference annotation *S*_*ann*_ for each frame. The expert annotations of this independent test set demonstrated high inter-rater agreement, with a Randolph’s Kappa [31] of 0.91±0.09 (range: 0.70-0.97). Detailed Randolph’s Kappa values for all TTE recordings are shown in Supplemental Figure 1 and Table 1. This benchmark dataset served as the reference standard for evaluating workflow performance, with automated measurements validated against expert-derived annotations.

**Table 1:** The table lists mouse and corresponding human gene abbreviations for the 37 genes identified by clustering analysis as deviating from the normal population. The physiological state during measurement is indicated (awk, awake; iso, anesthetized under isoflurane). Crossed boxes 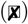 mark a link to mouse or human heart disease, based on a systematic in silico review, including genome-wide association study (GWAS) data. Hollow boxes indicate no reported association.

| Gene | Ortholog | State | Heart disease |  | Gene | Ortholog | State | Heart disease |  |
| --- | --- | --- | --- | --- | --- | --- | --- | --- | --- |
|  |  |  | Mouse | Human |  |  |  | Mouse | Human |
| *Acot12* | ACOT12 | awk | □ | □ | *Ndufv2* | NDUFV2 | iso | ✗ | ✗ |
| *Ankrd50* | ANKRD50 | iso | □ | □ | *Nupr1* | NUPR1 | iso | ✗ | ✗ |
| *Arhgap26* | ARHGAP26 | awk | ✗ | ✗ | *Odad3* | ODAD3 | awk | ✗ | ✗ |
| *Arhgef40* | ARHGEF40 | iso | □ | ✗ | *Orai3* | ORAI3 | iso | ✗ | ✗ |
| *Atp6v1a* | ATP6V1A | iso | □ | ✗ | *Pdcd6ip* | PDCD6IP | awk | ✗ | ✗ |
| *Atp8b3* | ATP8B3 | awk | □ | □ | *Pdgfc* | PDGFC | awk | ✗ | □ |
| *Car3* | CA3 | awk | ✗ | ✗ | *Pigk* | PIGK | iso | □ | □ |
| *Cdc45* | CDC45 | iso | □ | ✗ | *Pkm* | PKM | iso | ✗ | ✗ |
| *Cenpt* | CENPT | iso | □ | □ | *Rbm39* | RBM39 | awk | □ | □ |
| *Cep70* | CEP70 | iso | □ | ✗ | *Rpusd3* | RPUSD3 | iso | □ | □ |
| *Ckap2l* | CKAP2L | iso | □ | ✗ | *Sash1* | SASH1 | iso | □ | ✗ |
| *Ctc1* | CTC1 | awk | □ | ✗ | *Serpina3l* | SERPINA3 | iso | ✗ | ✗ |
| *Eea1* | EEA1 | awk | □ | □ | *Trio* | TRIO | iso | □ | ✗ |
| *Fam216a* | FAM216B | iso | □ | ✗ | *Tspan15* | TSPAN15 | awk | □ | □ |
| *Glmn* | GLMN | iso | ✗ | □ | *Uqcrh* | UQCRH | awk | ✗ | ✗ |
| *Golt1b* | GOLT1B | awk | ✗ | □ | *Urb1* | URB1 | iso | □ | □ |
| *Kctd2* | KCTD2 | awk | □ | □ | *Vdac1* | VDAC1 | awk | ✗ | ✗ |
| *Mrpl41* | MRPL41 | iso | □ | □ | *Zfp280d* | ZNF280D | awk | ✗ | □ |
| *Mybpc3* | MYBPC3 | awk | ✗ | ✗ |  |  |  |  |  |

### Validated segmentation model achieves high performance

Model performance was assessed using a 5-fold cross-validation procedure with distinct and non-overlapping training (80%) and validation (20%) subsets. Subsequently, five models *M*_*a*−*e*_ were trained under identical hyperparameter settings to evaluate consistency across folds. All models *M*_*a*−*e*_ were evaluated on the benchmarking dataset and demonstrated highly consistent segmentation performance, achieving a mean weighted Dice score of 97.60% (range: 97.54-97.64%) and a mean binary Dice score of 86.15% (range: 86.02-86.21%).

An additional key performance indicator was the signed difference between the predicted left ventricle inner diameter *S*_*pred*_ and the corresponding measurement derived from the expert-annotated reference mask *S*_*ann*_. The models *M*_*a*−*e*_ showed a mean signed deviation of -0.05 ± 0.20 mm relative to the expert annotation *S*_*ann*_, indicating a minimal underestimation of left ventricle inner diameter between model- and human-derived values. This deviation is considered negligible, given the inherent variability in manual border delineation that occurs even among expert human annotators due to image resolution (1 pixel = 0.02 mm), ambiguous boundaries, and subjective interpretation.

The best-performing model, *M*_*c*_, achieved a mean signed deviation of 0.04 ± 0.19 mm (2 pixels ± 9.5 pixels), a binary Dice score of 86.21%, and a weighted Dice score of 97.60%, and was therefore selected for all subsequent large-scale analyses. This level of model performance provides a robust foundation for reliable and biologically meaningful downstream analyses of TTE data.

### Automated quantification of cardiac parameters

The best-performing model, *M*_*c*_, was applied to the complete dataset to enable large-scale extraction of cardiac measurements. In total, 597,984,823 left ventricle inner diameters were quantified from the predicted segmentation masks *S*_*pred*_, achieving a mean confidence score of 0.01 ±0.01, where a value of 0 denotes maximal and 1 a minimal confidence in prediction.

Automated peak detection identified 1,853,687 systolic left ventricle inner diameters (*LVID*_*s*_) and 1,858,614 diastolic (*LVID*_*d*_) instances. Based on these inner diameters, key functional parameters, including end-systolic and end-diastolic volumes (*Vol*_*s*_, *Vol*_*d*_), heart rate (*HR*), stroke volume (*SV*), cardiac output (*CO*), fractional shortening (*FS*), and ejection fraction (*EF*) were calculated.

These comprehensive measurements across our large-scale dataset provided the foundation for establishing reference ranges in wild-type control mice and for subsequent detection of genotype-phenotype deviations in cardiac morphology and function across knockout lines present in the dataset.

### Control cohorts define normal cardiac morphology and function

Single-parameter reference ranges were established using 6,710 control mice, a subset of the 18,402 mice in the data set. We stratified their data by age (EA, early adult, mean age: 12 weeks; LA, late adult, mean age: 63 weeks), physiological state (awake and isoflurane anesthetized), and sex (female and male) (Figure 2). Due to the non-Gaussian distribution of the nine cardiac parameters, the median, along with the 2.5th and the 97.5th percentiles, was used to define the 95% reference ranges. No statistical comparative analyses were performed, as reference values served only to support visual inspection.

**Figure 1.**
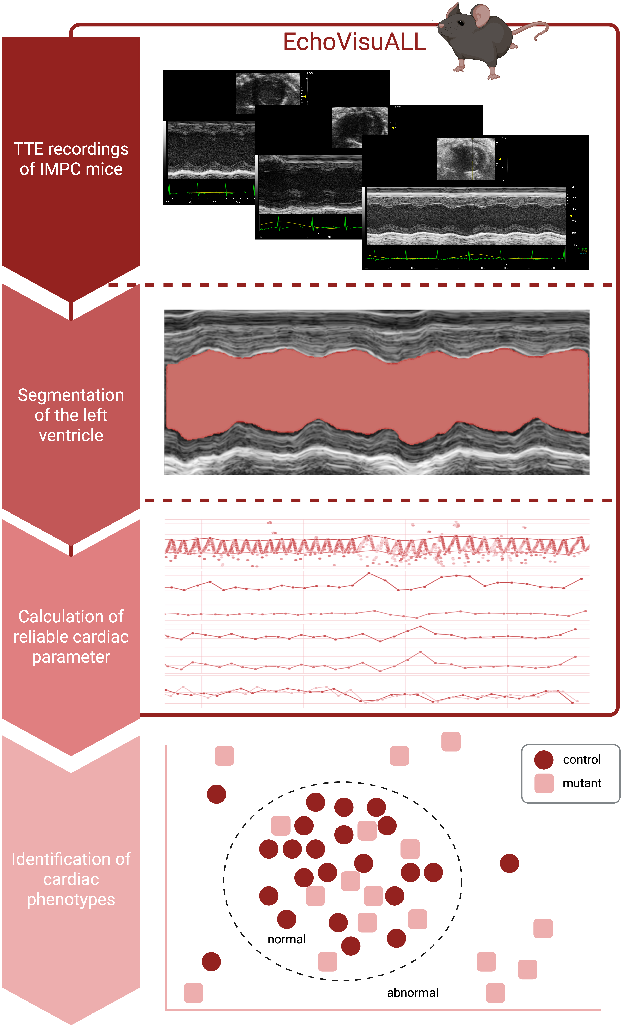
Graphical Abstract

**Figure 2.**
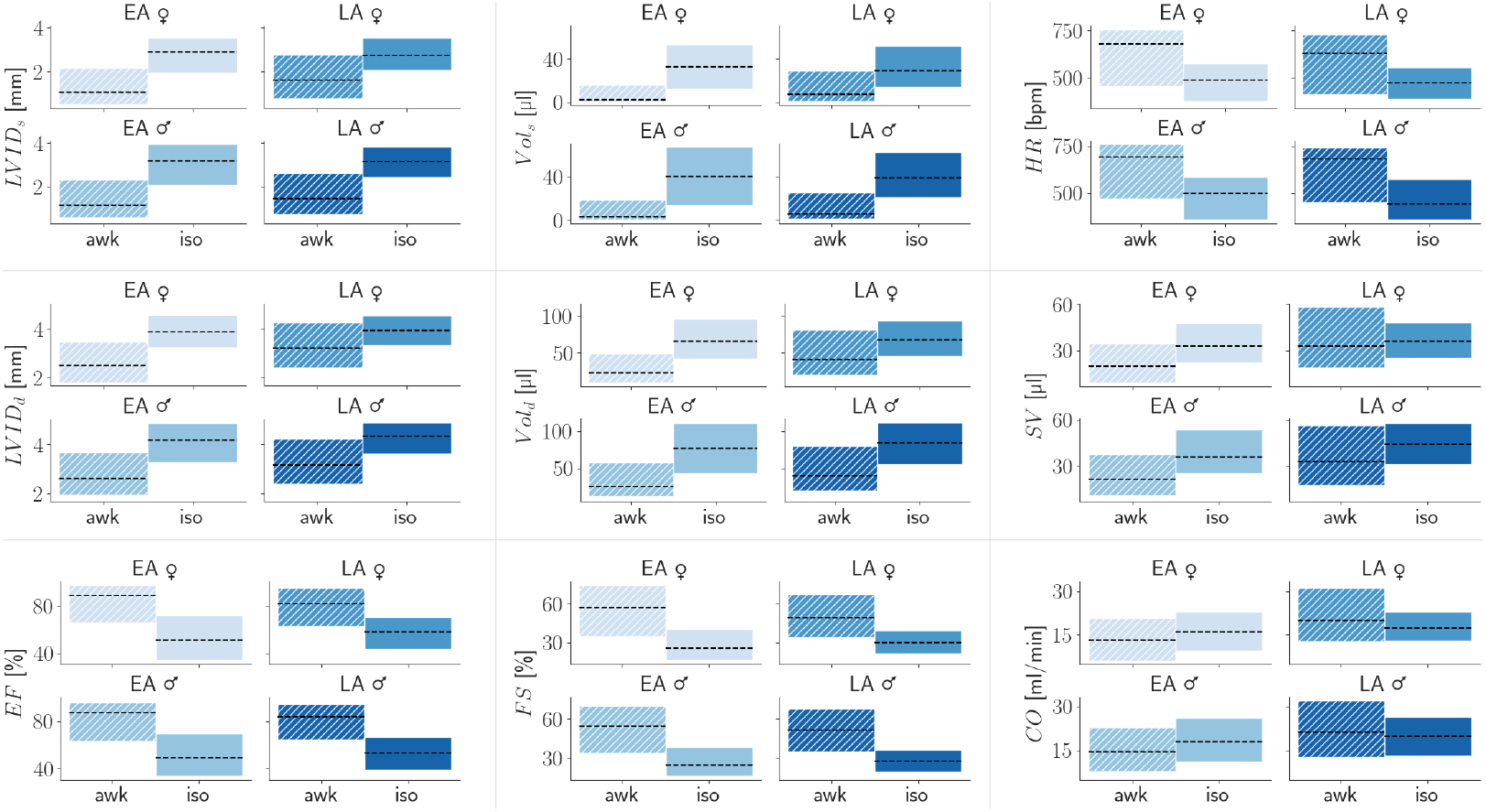
Control cohorts define normal cardiac morphology and function. Reference ranges for nine functional parameters were generated on 15,950 EA (early adult, mean age 12 weeks) and 2,430 LA (late adult, mean age 63 weeks) mice, stratified by age, physiological state, and sex. The boxes show the 2.5th and the 97.5th percentiles of the data distribution, while the dashed line indicates the median value within each reference range.

Distinct differences were observed between the two physiological states and ages consistent with previously reported TTE reference ranges [32]. Mice under isoflurane anesthesia exhibited increased *LVID*_*s*_, *Vol*_*s*_, *LVID*_*d*_, *Vol*_*d*_, and *SV*, along with decreased *HR, EF*, and *FS*, while *CO* remained comparable between groups. Moreover, late adult mice (LA, mean age: 63 weeks) showed larger *LVID*_*s*_, *Vol*_*s*_, *LVID*_*d*_, and *Vol*_*d*_ compared to early adult mice (EA, mean age: 12 weeks) in agreement with [32]. Comprehensive numeric reference ranges are provided in the Supplemental Table 2.

### Exploring cardiac phenotypes in mutant cohorts

Using the established reference ranges as a baseline, the mutant dataset comprised 11,692 mice, representing 715 distinct genes, each generated through single-gene knockout lines. Most gene knockout cohorts were homozygous (60%), while heterozygous (39%) and hemizygous (1%) mice were also included in the analysis. Cardiac parameters derived from the mutant cohorts exhibited overall distributions comparable to those of controls, with moderate variation across both morphological and functional measures provided in Supplemental Table 2. This variability likely reflects subtle gene-specific influences on the heart within the normal physiological spectrum. The resulting phenotypic heterogeneity provided the basis for subsequent data-driven clustering analysis aimed at identifying distinct cardiac phenotypes and uncovering genotype-phenotype associations.

### Data-driven clustering captures cardiac diversity

To identify inherent structure within the multidimensional space (9 cardiac parameters and their corresponding coefficients of variation) from mutant and control mice, a density-based clustering approach (DBSCAN [33]) was applied to detect phenotypically abnormal individual mice relative to the reference cohort. Multidimensional cluster analyses were performed separately for sex (females and males), physiological state (awake and anesthetized), and age (EA and LA), resulting in eight distinct subgroup analyses.

To ensure robustness and stability of cluster detection, the DBSCAN *epsilon* hyperparameter was systematically modified across five categories I-V (Table 4). This iterative adjustment allowed discrimination between mice exhibiting a strong and well-defined phenotype (category I) and those showing progressively weaker deviations (category V). The refinement was finished when the normal cluster’s composition indicated a decreasing proportion of control mice. The applied *epsilon* values ranged from 1.9-1.5 for EA-anesthetized, 1.7-1.3 for EA-awake, 2.0-1.6 for LA-anesthetized, and 1.8-1.4 for LA-awake mice. Importantly, sex did not affect the *epsilon* ranges.

Collectively, this five-category procedure identified 37 of 715 genes with distinct phenotypic structure in the multidimensional cardiac parameter space. These genes were subsequently analyzed in greater detail to validate the multivariate approach and to identify gene-specific cardiac phenotypes.

### Identification of genes associated with heart disease

The density-based clustering analysis identified 37 genes for which at least one mutant cohort with a size of 4–16 mice, differing in age, physiological state, sex, and zygosity, was classified as phenotypically abnormal. A cohort was considered abnormal when at least 75% of its constituent mice were identified as outliers and hereby categorized as abnormal within the clustering framework. To confirm these findings, the 9 cardiac parameters of each individual mouse were compared against the corresponding reference ranges established for the respective subgroup, thereby supporting all identified phenotypic deviations. In addition, all related but not phenotypically abnormal mutant cohorts were systematically examined and compared with their reference distributions to obtain a comprehensive characterization for each of the 37 genes and their respective cardiac phenotypes.

A complete list of all identified genes is provided in Table 1, in alphabetical order. Of these, 17 genes were identified in cohorts of awake mice, and 20 genes in cohorts under isoflurane. Among the abnormal cases, 24 genes were detected exclusively in the EA subset and 11 in the LA subset, whereas 2 genes were phenotypically abnormal in both age groups (*Mybpc*3 and *Zfp*280*d*). A sex-specific pattern was observed, with 22 genes classified as abnormal in males, 11 in females, and 4 showing abnormalities in both sexes.

Beyond identifying genes associated with abnormal cardiac phenotypes, the multivariate clustering framework additionally produced a ranked list of genes, ordered according to the magnitude of phenotypic deviations from the corresponding reference control population.

### Top-ranked genes reveal distinct cardiac phenotypes

The ranking procedure identified 11 top-ranked genes (Table 2) that consistently exhibited the most robust and pronounced cardiac phenotypes across different clustering configurations. These genes were categorized as abnormal under the largest epsilon (I, Table. 4) applied within each subgroup (EA-anesthetized: 1.9; EA-awake: 1.7; LA-anesthetized: 2.0; EA-awake: 1.8).

**Table 2:** The table summarizes the 11 top-ranking genes identified by clustering analysis. Information includes mouse gene name, cohort characteristics—early adult (EA, median age: 12 weeks) and late adult (LA, median age: 63 weeks)—as well as zygosity and sex. The corresponding human ortholog and full gene name are also provided. Abnormal cohorts were marked with a crossed box 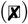; hollow boxes indicate cohorts not classified as abnormal, and a dash (–) denotes missing or unavailable cohorts.

| Gene | EA |  |  |  | LA |  | Ortholog, gene name |
| --- | --- | --- | --- | --- | --- | --- | --- |
|  | ♀<br>Hom | Het | ♂<br>Hom | Het | ♀<br>Hom | ♂<br>Hom |  |
| *Acot12* | □ | - | ⊠ | - | - | - | ACOT12, acyl-CoA thioesterase 12 |
| *Atp8b3* | - | □ | - | ⊠ | - | - | ATP8B3, ATPase, class I, type 8B, member 3 |
| *Cep70* | ⊠ | - | ⊠ | - | - | - | CEP70, centrosomal protein 70 |
| *Eea1* | ⊠ | □ | ⊠ | □ | - | - | EEA1, early endosome antigen 1 |
| *Kctd2* | □ | - | ⊠ | - | - | - | KCTD2, potassium channel tetramerisation domain containing 2 |
| *Mybpc3* | ⊠ | □ | ⊠ | □ | ⊠ | ⊠ | MYBPC3, myosin binding protein C, cardiac |
| *Nupr1* | - | - | - | - | □ | ⊠ | NUPR1, nuclear protein transcription regulator 1 |
| *Odad3* | - | ⊠ | - | □ | - | - | ODAD3, outer dynein arm docking complex subunit 3 |
| *Tspan15* | ⊠ | □ | □ | □ | - | - | TSPAN15, tetraspanin 15 |
| *Vdac1* | - | □ | ⊠ | □ | - | - | VDAC1, voltage-dependent anion channel 1 |
| *Zfp280d* | □ | - | ⊠ | - | □ | ⊠ | ZNF280D, zinc finger protein 280D |

For comprehensive phenotypic characterization, all mutant cohorts associated with these genes, including those not categorized as abnormal, were analyzed to capture the full range of cardiac variation. In total, 17 mutant cohorts showed measurable deviations in one or more cardiac parameters. The majority of these deviations occurred in male homozygous cohorts (7/17 in EA and 3/17 in LA mice). Among female homozygous cohorts, 4/17 were identified in EA and 1/17 in LA groups. In instances where both homozygous and heterozygous cohorts were available, heterozygous mice consistently lacked substantial phenotypic abnormalities, a finding consistent with biological expectations and supporting the discriminative power of the clustering framework.

Notably, for *Atp*8*b*3 and *Odad*3, phenotypic alterations were sufficiently pronounced to be detected even in heterozygous cohorts, while homozygous mice were reported in the IMPC to have preweaning lethality with incomplete penetrance (www.mousephenotype.org). The results for *Atp*8*b*3 and *Odad*3 are biologically plausible, since lethality of homozygous animals is consistent with a strong heterozygous phenotype. Among the genes assessed in both age groups, *Mybpc*3 and *Zfp*280*d* exhibited concordant phenotypic deviations in EA and LA cohorts, suggesting persistence or early onset of cardiac alterations.

Although these 11 genes displayed the most consistent and biologically plausible phenotypes within the multivariate clustering analysis, each gene candidate underwent an extensive literature review to confirm concordance with previously reported cardiac phenotypes (Supplemental Table 3).

### Categorization of genes based on cardiac association and GWAS evidence

To complement the computational findings and assess known disease associations, a systematic literature review was conducted for all 37 genes identified through multivariate clustering. This in silico survey aimed to determine previously reported links between these genes and cardiac pathophysiology, without restricting the search to the specific phenotypes identified in this study. The review revealed distinct gene categories, separating those with established cardiac associations from genes with no prior connection to the heart.

Among the 37 genes, 25 were categorized as “proof-of-concept genes”, having previously been associated with heart disease or related traits in humans or mice, including genes supported by genome-wide association study (GWAS) evidence. Specifically, 8 of them are linked to human heart disease, 4 to mouse models and 13 to both species. The remaining 12 genes represent “novel candidate genes” with no prior association to heart function or pathology (Table 1). A comprehensive overview of all identified genes, associated mouse cohort metadata, and corresponding literature references is provided in the Supplemental Table 3.

To illustrate these categories, two proof-of-concept genes (*Mybpc*3 and *Cep*70), and one novel candidate (*Acot*12) were selected as representative examples for detailed characterization in the following sections.

### *Mybpc3*: A candidate gene for multivariate cardiac analysis

Comprehensive phenotyping of Myosin-binding protein C, cardiac-type (*Mybpc*3) mutant mice revealed pronounced alterations in cardiac morphology and systolic function across age, sex, and zygosity. Cohorts of male and female mice were analyzed at early adult (EA, mean age: 12 weeks) and late adult (LA, mean age: 63 weeks) stages, with all echocardiographic measurements performed in awake mice. The EA cohort included both homozygous and heterozygous genotypes, enabling a detailed assessment of the gene dosage effect (Figure 3 Panel b). Multivariate cluster analysis, validated by parameter-specific reference ranges, uncovered a consistent pattern of systolic impairment in *Mybpc*3 deficient mice.

**Figure 3.**
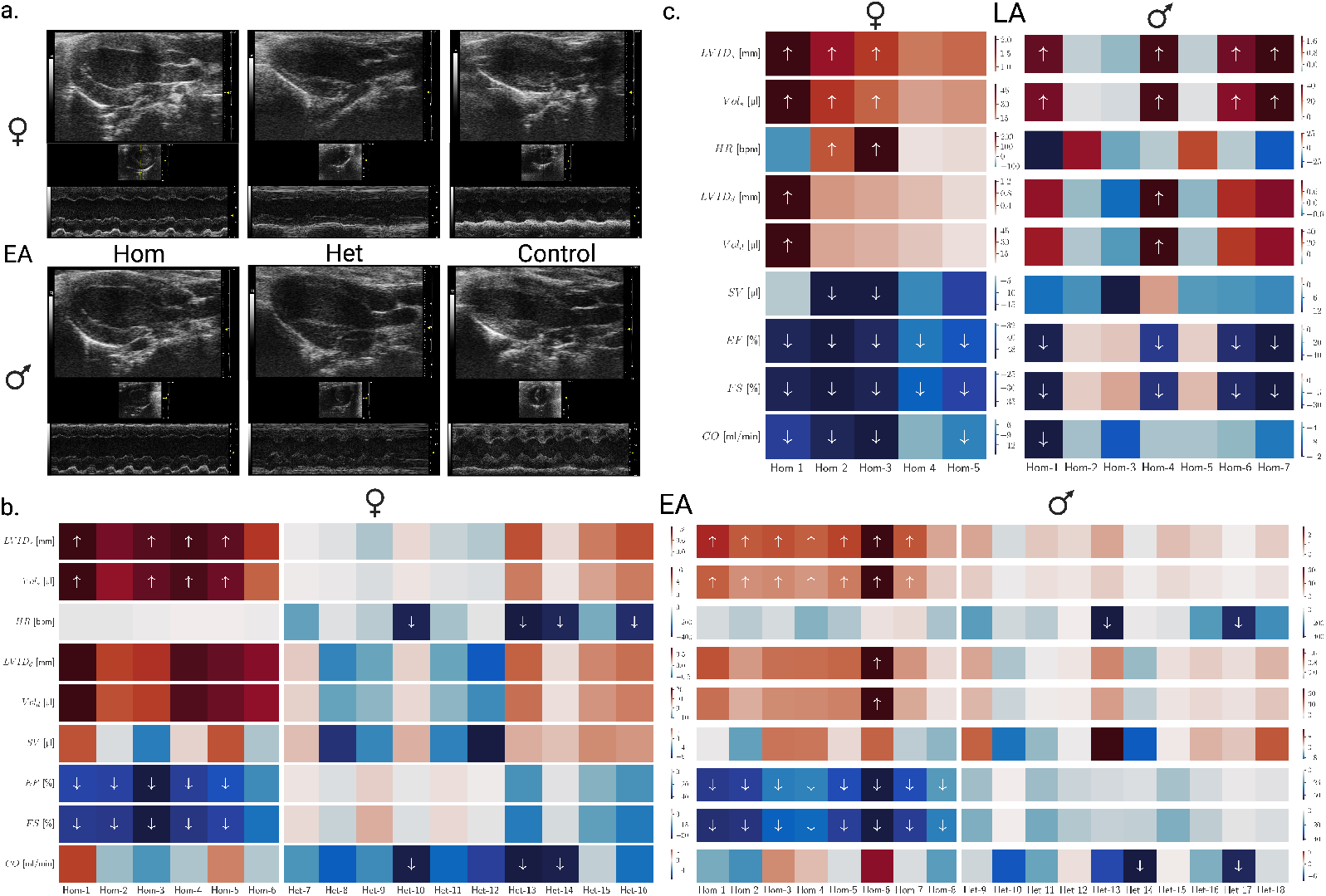
Homozygous *Mybpc3* mutants of both sexes displayed ventricular dilation and reduced systolic performance (*LV ID*_*s*_, *V ol*_*s*_, ↓*EF*, ↓*FS*), while heterozygotes largely remained normal. The phenotype persisted at later age with moderate attenuation, confirming *Mybpc3* deficiency as a robust determinant of systolic dysfunction. Representative parasternal long- and short-axis echocardiography recordings illustrate characteristic phenotypes across zygosities in EA (panel a). Heatmaps show single-parameter deviations for homozygous and heterozygous *Mybpc3* cohorts stratified by sex and age (EA, 12 weeks, panel b; LA, 63 weeks, panel c). The arrows indicate the direction of abnormal single-parameter changes.

In EA female mice (n=16; 6 homozygous and 10 heterozygous), all homozygous individuals were categorized as abnormal, with 4/6 showing increased *LVID*_*s*_ and *Vol*_*s*_, and 5/6 exhibiting reduced *EF* and *FS*. EA male mice (n=18; 8 homozygous, 10 heterozygous) displayed a comparable phenotype. All homozygous males were identified as abnormal, 7/8 showing ventricular dilation (increased *LV ID*_*s*_ and *Vol*_*s*_), and all showed markedly reduced *EF* and *FS*. Heterozygous mice of both sexes largely remained within normal limits.

At the later age, the phenotype persisted. All LA homozygous females (n=5) exhibited systolic abnormalities with 3/5 showing increased *LVID*_*s*_ and *Vol*_*s*_, 5/5 reduced *EF* and *FS*, and 4/5 decreased *CO*. In LA homozygous males (n=7), 6/7 mice were categorized as abnormal, with 4/7 demonstrating ventricular dilation (increased *LVID*_*s*_ and *V ol*_*s*_) and reduced systolic performance (decreased *EF* and *FS*) with less pronounced changes than those observed in younger cohorts. Collectively, these findings establish *Mybpc*3 deficiency as a strong determinant of systolic dysfunction characterized by ventricular dilation and impaired contractility across age and sex, illustrated in Figure 3.

*Mybpc*3 is a major sarcomeric component with both structural and regulatory functions [34]. Consistent with the penetrance of its phenotype (homozygous vs. heterozygous) and its identification as a candidate gene in our clustering analysis, it was used as a proof-of-concept gene to validate the robustness and accuracy of our multivariate clustering and reference-range approach.

### Functional evidence for cardiac involvement of *Cep70*

For the phenotypic characterization of centrosomal protein 70 (*Cep70*), measurements were obtained from 8 EA mice (n=4 females, n=4 males), all homozygous for the targeted allele and examined under isoflurane anesthesia. Multivariate analysis revealed a distinct cardiac phenotype characterized by smaller ventricular dimensions, elevated heart rate, and higher systolic indices compared with reference ranges, most prominently in females.

The clustering analysis detected all female mice were as abnormal, whereas 3/4 of the male mice were identified as abnormal. Single-parameter evaluation partially supported these findings: Among females, 2/4 exhibited a consistent and distinct phenotype showing reduced *LVID*_*s*_, *LVID*_*d*_ and *Vol*_*s*_, *Vol*_*d*_, along with decreased *SV* and increased *HR, EF*, and *FS*, while the remaining two did not differ from reference values. In males, 3/4 animals showed a phenotype similar to that of the affected females, characterized by reduced *LVID*_*d*_, *Vol*_*d*_, and *SV* . One male exhibited a reduction in *SV, HR*, and *CO* relative to the reference ranges, whereas another showed no alterations except for a decrease in *SV* .

Collectively, these findings indicate a small-cavity cardiac phenotype in a subset of mice with relatively higher measures of systolic function and heart rate compared with reference values, suggestive of an enhanced cardiac performance state, as illustrated in Figure 4. *Cep70* has also been linked through GWAS studies to coronary artery disease, myocardial infarction, septal mass, and isolated congenital heart disease [35, 36, 37], yet the present experimental mouse data are valuable for investigating the cardiac impact of *Cep70*.

**Figure 4.**
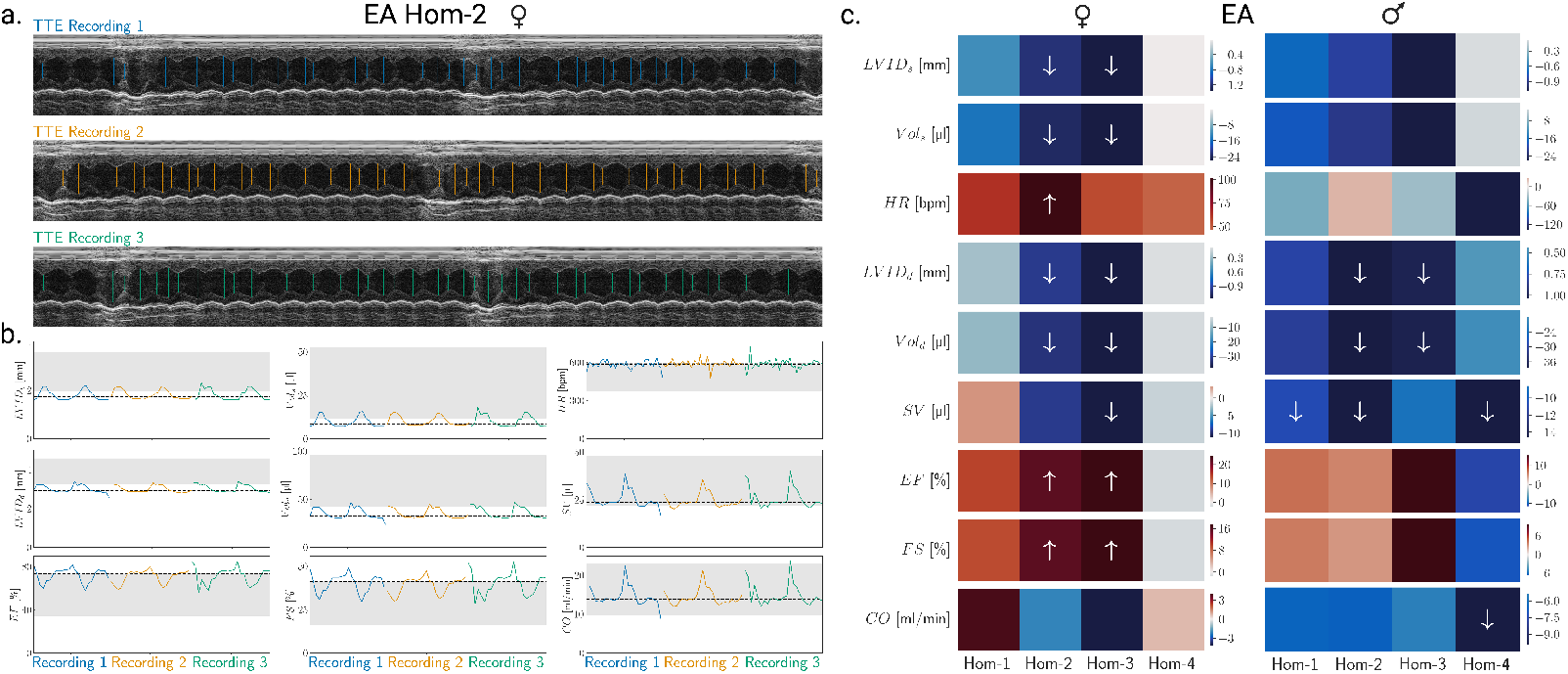
All *Cep70* homozygous females and most males exhibited reduced ventricular size and increased systolic indices. A representative *Cep70* female (Hom-2) shows parasternal short-axis M-mode echocardiography recordings with *LV ID*_*s*_, *LV ID*_*d*_ indicated as min and max across the entire recording (panel a). Corresponding parameter values of *LV ID*_*s*_, *V ol*_*s*_, *HR, LV ID*_*d*_, *V ol*_*d*_, *EF, FS*, and *CO* were plotted against grey reference ranges (panel b). Cohort-level results from multivariate clustering and single-parameter analyses, stratified by sex, are shown in panel c. The arrows indicate the direction of abnormal single-parameter changes. Abnormal profiles with single-parameter validation confirm abnormalities in 2/4 females and 4/4 males in the young adult stage (EA).

### *Acot12* as a potential new candidate for dilated cardiomyopathy

In *Acot12*, Acyl-CoA Thioesterase 12, transthoracic echocardiographic measurements were obtained from 19 awake homozygous mice (n=10 females and n=9 males). Within the clustering analysis, 7/9 male mice were identified as abnormal, whereas only 4/10 females were categorized as abnormal. Consequently, the female cohort did not exceed the 75% threshold and was therefore not labeled as abnormal.

In males, 6/9 exhibited a distinct dilated phenotype, characterized by enlarged ventricular dimensions (*LVID*_*s*_, *LVID*_*d*_) and volumes (*Vol*_*s*_, *Vol*_*d*_, *SV*) together with reduced *EF, FS*, and *HR*. The remaining three male mutants showed subtle or no deviations from the reference ranges. In females, 6/10 exhibited comparable but less pronounced alterations, remaining predominantly within the normal reference ranges. Collectively, these findings indicate a sex-dependent dilated cardiomyopathy-like phenotype characterized by ventricular enlargement and impaired systolic performance, with greater severity in males, as illustrated in Figure 5. To date, no association between *Acot12* and cardiac disease has been reported, and the mouse data presented here provide functional evidence for a previously unrecognized link to dilated cardiomyopathy, identifying *Acot12* as a potential novel candidate gene for this condition.

**Figure 5.**
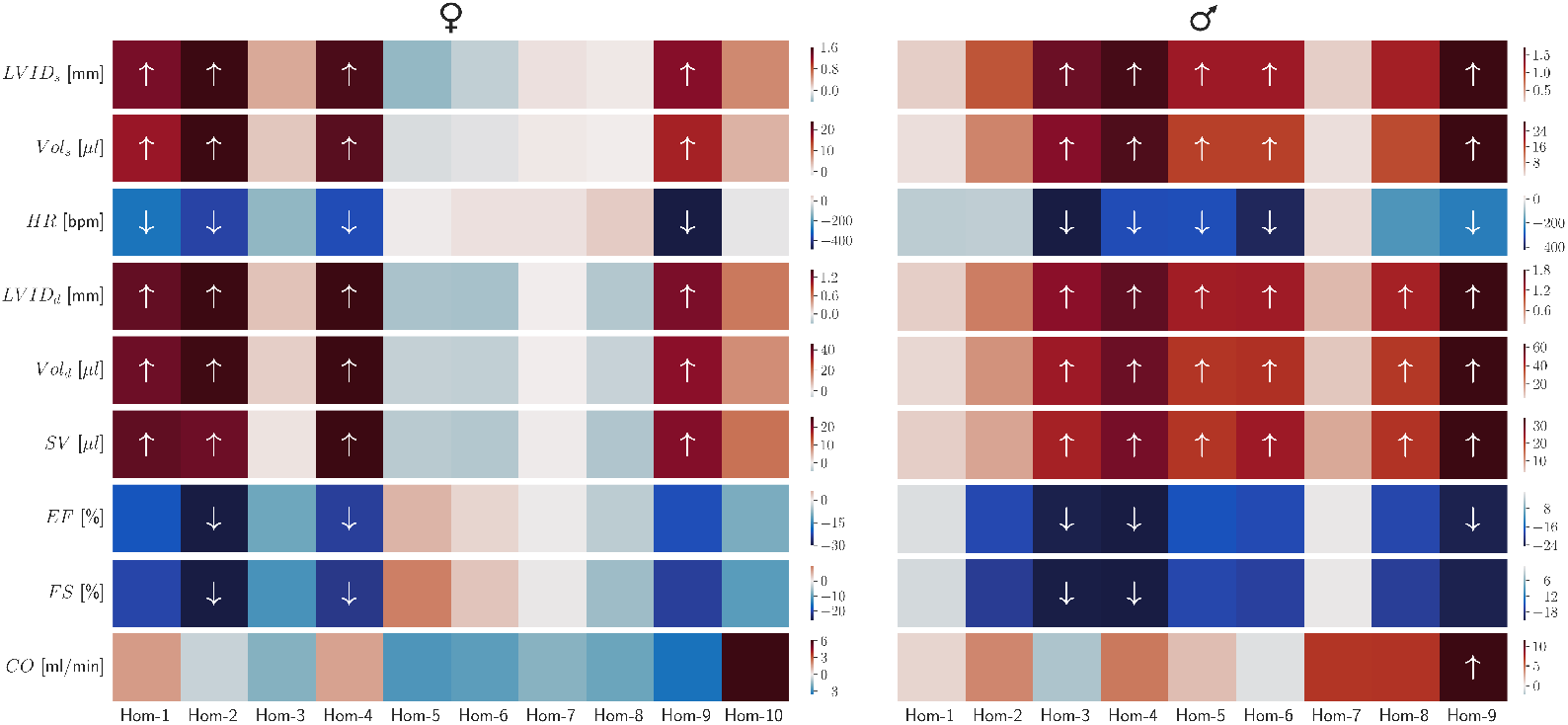
Males exhibited a dilated phenotype with enlarged ventricular dimensions and volumes and reduced *EF, FS*, and *HR*, consistent with male-predominant systolic dysfunction. Heatmaps show single-parameter deviations of awake early adult (EA) homozygous *Acot12* mice, stratified by sex. 7/9 males (right) and 4/10 females (left) were classified as abnormal. Arrows indicate the direction of abnormal single-parameter changes.

Together, *Mybpc3, Cep70* and *Acot12* exemplify key categories among the 37 genes identified by multivariate clustering and supported by comparisons with single-parameter reference ranges: a known cardiac gene (*Mybpc3*), a GWAS-derived candidate with new functional mouse evidence (*Cep70*), and a previously unlinked gene now associated with dilated cardiomyopathy (*Acot12*).

## Discussion

This study introduces an integrative framework for large-scale cardiac phenotyping, combining automated high-throughput transthoracic echocardiography (TTE) with multidimensional, unsupervised clustering. Unlike conventional imaging-based, genotype–phenotype studies that rely on selected metrics or manual interpretation, this approach leverages comprehensive image-derived data to uncover complex and previously unrecognized cardiac phenotypes. Analysis of 65,241 TTE recordings from 18,402 mice revealed 37 genes with substantial cardiac abnormalities, encompassing both known, potentially shared across the murine and human systems, and novel genetic associations. By combining large-scale image analysis with data-driven, multidimensional analysis, this work establishes a scalable paradigm for systematic, unbiased exploration of cardiac function. The integration of automated segmentation with unsupervised clustering not only enhances phenotypic resolution but also redefines how genetic determinants of cardiovascular physiology can be discovered and interpreted in a data-intensive context.

The automated image analysis pipeline presented here provides a robust foundation for unbiased, high-throughput quantification of cardiac morphology and function. By minimizing operator dependency and incorporating confidence metrics for segmentation reliability, EchoVisuALL ensures both scalability and reproducibility across large datasets. Beyond standardization, this approach overcomes a key limitation in cardiac phenotyping and addresses one of the long-standing challenges in large-scale cardiac phenotyping, capturing nonlinear relationships among parameters that may not be apparent in univariate analyses. Through multidimensional clustering, all parameters are evaluated in concert, yielding an integrated view of cardiac function rather than isolated metrics. Incorporating the coefficient of variation as a measure of temporal stability further introduces a dynamic perspective, capturing variability across cardiac cycles. Together, these elements create a data-driven framework that improves genotype-phenotype resolution and reveals subtle gene-specific effects on cardiac physiology.

The integration of automated cardiac phenotyping with multidimensional analysis provided a powerful platform to bridge experimental genetics with human cardiovascular biology. Among the 37 genes identified, several align closely with known human cardiac disease mechanisms, underscoring the translational potential of this approach. For example, *Mybpc3*, one of the most frequently mutated genes in hypertrophic cardiomyopathy [38, 39, 40, 41], displayed ventricular dilation and impaired contractility in knockout mice. This phenotype is consistent with a complete loss of sarcomeric integrity and represents an end-staged dilated cardiomyopathy [42, 43, 44], mirroring aspects of the human disease spectrum and validating the workflow’s capacity to recapitulate clinically relevant phenotypes.

In contrast, *Cep70* emerged as a less characterized but particularly intriguing gene. The mouse data revealed a small-cavity phenotype accompanied by elevated systolic function and increased heart rate, suggestive of a hypercontractile state and enhanced cardiac performance. Notably, *CEP70* has been linked through GWAS studies to coronary artery disease, myocardial infarction, septal mass and isolated congenital heart disease [35, 36, 37]. However, no experimental evidence has previously substantiated these associations. Our findings thus provide the first functional evidence connecting *Cep70* disruption to cardiac remodeling, establishing a foundation for targeted mechanistic studies in both animal models and human tissues.

Similarly, *Acot12*, which encodes a cytosolic acyl-CoA thioesterase involved in lipid metabolism [45], has not been previously associated with heart disease. The observed dilated cardiomyopathy phenotype in *Acot12*-deficient mice introduces a new link between myocardial energy homeostasis and structural remodeling, suggesting that metabolic dysregulation may contribute to cardiac dilation in previously unrecognized ways.

Remarkably, among the top-ranked genes identified through multivariate clustering, *Atp8b3, Eea1, Kctd2* and *Tspan15* emerged as novel candidate genes with robust and consistent cardiac phenotypes, despite no prior associations with cardiovascular function or pathology. Their discovery broadens the genetic landscape of cardiac biology, extending beyond canonical pathways. *ATP8B3*, a phospholipid flippase [46], could affect sarcolemmal membrane integrity and ion channel distribution, offering a potential mechanistic bridge between membrane remodeling and electrophysiological stability. *EEA1*, a key regulator of endosomal trafficking [47], points to a possible role of intracellular vesicular dynamics in cardiomyocyte homeostasis and stress adaptation. KCTD2, part of a protein family implicated in Cullin3-dependent ubiquitination [48], may influence myocardial protein turnover and the cellular response to hemodynamic or metabolic stress. Finally, *TSPAN15*, a member of the tetraspanin superfamily, may participate in cell–cell communication and extracellular matrix interactions that underpin cardiac remodeling [49].

In addition to these novel candidates, among the 25 established heart-associated genes (*Arhgab26, Arhgef40, Atp6v1a, Car3, Cdc45, Cep70, Ckap2l, Ctc1, Fam216a, Glmn, Golt1b, Mybpc3, Ndufv2, Nupr1, Odad3, Orai3, Pdcd6ip, Pdgfc, Pkm, Sash1, Serpina3l, Trio, Uqcrh, Vdac1, Zfp280d*), eight show compelling evidence for involvement in human cardiac disease but currently lack suitable experimental models. The mouse data presented herein may thus provide a valuable framework to investigate these genes in vivo, enabling the study of distinct and potentially underexplored cardiac pathologies.

Collectively, these findings exemplify how large-scale, data-driven cardiac phenotyping can both validate established human gene-disease relationships and reveal new candidates with translational potential. Equally important, the automated pipeline and the multidimensional clustering framework are universally applicable across mouse echocardiography datasets encompassing variation driven by age, sex, and anesthesia, enabling retrospective reanalysis of existing studies to validate manual findings, uncover hidden patterns and extract greater biological depth from archived data.

The human expert-curated, gold-standard dataset generated in this study serves as a benchmark for the field, providing a reliable reference for training and refining automated analysis tools such as EchoVisuALL. This resource also facilitates the calibration of algorithms applied to datasets outside the current analytical scope, supporting consistent and accurate quantification across a wide range of imaging conditions.

While our pipeline has facilitated a multidimensional approach to genotype-phenotype discovery, it still preserves and enhances the ability to generate and compare against single-parameter reference ranges at scale. This enables a rigorous validation of these cardiac metrics across mouse models, improving reproducibility and comparability between studies. Together, these advances establish a transparent framework for automated cardiac phenotyping, one that not only accelerates gene discovery in experimental mouse models but also lays the groundwork for translating gene information and imaging data into deeper insights into heart disease.

Despite its strengths, this study has several important limitations that must be considered. The framework presented here focuses exclusively on masking the left ventricular inner diameter and derives cardiac parameters using the Teichholz formula [50, 51]. As such, septal mass, wall thickness, and right heart dimensions or contractility, as well as blood flow metrics for diastolic function, were not integrated into the analysis. This limitation restricts the ability to fully assess biventricular and structural remodeling patterns that contribute to heart dysfunction.

The workflow was also not evaluated in interventional models such as myocardial infarction, pressure overload, or diet-induced interventions (e.g., high-fat feeding leading to metabolic disturbances), which may exhibit distinct pathophysiological trajectories. Functional assessment relied solely on echocardiographic parameters without complementary histological, electrophysiological, or molecular validation. Furthermore, variation in cohort sizes and sex distribution across mutant cohorts may have influenced the strength or specificity of genotype-phenotype associations within the multidimensional clustering. Like any other AI-based approach, performance depends on appropriate hyperparameter tuning and data quality, although systematic cross-validation across workflow components enhanced robustness and reproducibility. Finally, although reference ranges provided a consistent internal baseline for cardiac normality, they may not fully capture inter-experimental variability or strain-specific differences. Future work should expand the analytical framework to incorporate right-heart and structural parameters, integrate additional phenotyping data, and validate findings in interventional and longitudinal models to broaden translational applicability.

Taken together, EchoVisuALL provides a robust and accessible framework for large-scale, multi-center, automated cardiac phenotyping, uniting image analysis and data integration into a scalable pipeline. By combining quantitative echocardiographic characterization with multidimensional clustering, it both validates known human gene-disease associations and uncovers novel genetic candidates with translational potential.

Looking ahead, the framework presented here can be extended to interventional models, such as myocardial infarction, pressure overload, or diet-induced metabolic disturbances, and integrated with molecular, electrophysiological, and histological data to enable truly multimodal insights. The resulting benchmark dataset provides a valuable foundation for retraining, adaptation to diverse TTE data and cross-species model development. Through transfer learning and cross-species validation, EchoVisuALL has the potential to bridge experimental genetics and cardiovascular medicine, advancing precision phenotyping and accelerating discovery in heart disease research.

## Methods

### The International Mouse Phenotyping Consortium

The International Mouse Phenotyping Consortium (IMPC) is a global research initiative with 24 institutional members [10]. Its aim is to generate single-gene knockouts (KO) for all protein-coding genes and to comprehensively characterize their phenotypes across a wide spectrum of organs based on standardized IMPReSS protocols [52, 53]. To achieve consistency across all participant centers, animal experiments, housing, and husbandry follow standardized procedures in compliance with the ARRIVE guidelines [54], ensuring both animal welfare and reproducibility. Further details are available on the IMPC portal (http://www.mousephenotype.org/about-impc/arrive-guidelines).

### IMPC Centers Contributing Transthoracic Echocardiogram Data

In this study, we used transthoracic echocardiogram (TTE) recordings that are not accessible via the IMPC portal from a subset of five IMPC data-contributing centers (ethical approval details are included in parentheses after each center):

- Baylor College of Medicine, BCM (Institutional Animal Care and Use Committee approved license AN-5896)
- Czech Center for Phenogenomics, CCP (AV CR 62/2016, Academy of Sci., Czech Rep.)
- Helmholtz Munich, Institute of Experimental Genetics and German Mouse Clinic, HMGU (#144-10, 15-168)
- Institute Clinique de la Souris, ICS (#4789-2016040511578546v2)
- Center for Phenogenomics, TCP (28-0045H).

### Animals

In total, 18,402 mice, both males and females, were included in this study, each with corresponding TTE recordings. These were acquired over 11 years from 2013 to 2024 and comprise acquisitions from 6,710 (36%) wild-type control and 11,692 (64%) mutant mice. Mutant mice were mainly CRISPR/Cas-based [52] and derived from inbred substrains of a C57BL/6N genetic background: C57BL/6NCrl (CCP, HMGU and ICS); C57BL/6NJ (BCM) and C57BL/6NTac (HMGU, ICS). Gene selection was not biased towards heart dysfunction. TTE recordings were collected from mice at one of two possible time points. A total of 87% (15,970 mice) of the data were collected in early adult (EA) age with a mean age of 12 weeks (minimum 8, maximum 16 weeks). The remaining 13% (2,432 mice) were collected in the late adult (LA) age, with a mean of 63 weeks (minimum 51, maximum 78 weeks).

### Transthoracic Echocardiogram (TTE) recordings

TTE was performed on awake mice or mice anesthetized with isoflurane, based on IMPC standard operating procedure (https://www.mousephenotype.org/impress/ProcedureInfo?procID=109). In brief, body weights were taken shortly before TTE recording. For anesthetized TTE recordings, the animal was placed in an induction chamber and anesthetized with 1.5-3% isoflurane. While sedated, either as part of the TTE session or as a separate preparatory procedure, the animal undergoes hair removal of the chest. With the hair removed, the animal was placed on the imaging platform with its paws taped to ECG surface electrodes and a rectal probe inserted to monitor body temperature, which was maintained at 36-37°C. During imaging, anesthesia was adjusted to maintain proper heart rate and keep the animal from waking up [55].

For awake TTE examinations, the animal was firmly held by the nape (in the supine position) in the palm of one hand, with the tail held tightly between the last two fingers. To facilitate ultrasound imaging, pre-warmed ultrasound gel was placed on the chest at the area of imaging, and TTE recordings were captured. For the short-axis mode, papillary muscles were used as an anatomic point of reference. For the parasternal long-axis, the apex and aortic root were used as anatomic points of reference. At least three TTE recordings in motion-mode (M-Mode) were captured for each mouse. Once imaging was complete, the animal was removed from the platform and allowed to recover atop a heating pad [55].

### Technical characteristics of TTE recordings

All TTE recordings were generated with the Vevo2100 or Vevo3100 (FUJIFILM Visualsonics, Inc.) imaging systems. The imaging frequencies varied among 24, 30, 32, and 40 MHz with a mean focus depth of 0.65 ± 0.68. All echocardiograms were exported in a DICOM format that includes multiple frames and uses RGB color encoding (3 channels per pixel). On average, each DICOM file contains 42±13 frames, with a frame width of 1022±30*px* and a frame height of 344±4*px*. The pixel resolution of the y-axis is 0.02 ± 0.02*mm/px*, and the time resolution of the x-axis 0.8 ± 0.02*ms/px*.

### Training data of the segmentation network

During an active learning process, 1,000 frames were annotated by human experts as detailed in [28]. Across multiple iterations, the most challenging frames were progressively selected and annotated. Based on quality, experts annotated either the entire frame or just specific sections. In this study, the pre-annotated 1,000 frames were quality-controlled and used to train a Bayesian U-Net. The frames were split 80/20 into training and validation data. In detail, the 1,000 frames were distributed as follows: 880 frames (88%) derived from EA and 120 frames (12%) from LA age; 258 frames (26%) from anesthetized and 742 frames (74%) from awake mice; 402 frames (40%) from control and 598 frames (60%) from mutant mice.

### Expert-curated benchmarking data

To evaluate the performance and generalization of our model, we created an independent validation dataset. This dataset contains 20 TTE recordings not previously seen by the model. They were randomly selected from four IMPC centers (BCM, ICS, HMGU, CCP), five per center. Prior to manual annotation, the recordings were cropped into 836 frames, each with a size of 1024×344 pixels. Additional metadata was removed to ensure unbiased annotation.

Next, each frame was independently annotated by five different IMPC experts using the VGG Image Annotator [56]. The five expert-based annotation masks, denoted as *a*_1-5_, were aggregated to one single segmentation mask 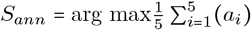, following [28]. As binary segmentation masks were used, this calculation is equivalent to a majority vote. Consequently, a pixel was classified as background if at least three annotations labeled it as 0. Conversely, a pixel was classified as part of the left ventricle if at least three annotations labeled it as 1. Finally, Randolph’s Kappa [31] was computed for each pixel to assess inter-rater agreement.

### Training of a Bayesian U-Net

We trained a Bayesian U-Net to generate a binary segmentation mask that labels the left ventricle in individual frames across a TTE recording. A total of 1,000 manually annotated frames served as the training dataset. The general setup was adapted as described in [28], including the model architecture of a U-Net [57] with dropout layers, the optimizer Adam [58], and a combined loss function that consists of binary cross-entropy and Dice loss [59]. The network uses a sigmoid activation function, a dropout rate of 0.2, a batch size of 1, and an input size of 1024×1024×3 pixels. To improve the performance and generalization of our model, data augmentation techniques (random brightness and contrast) were incorporated. The number of training epochs was increased to 200 with early stopping after 150 epochs to prevent overfitting. A learning rate scheduler decreased the learning rate from 0.0001 to 0.00001 after 100 epochs, and to 0.000001 after 150 epochs. For evaluation, a 5-fold cross-validation was performed to ensure a balanced training dataset and stable model performance.

### Confidence score

To assess the model’s confidence, a pixel-wise confidence score was computed [28]. Let 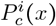 denote the probability for a given pixel *x* belonging to class *c* ∈ {*background, leftventricle}*. With *N* being the number of Monte Carlo samples and *i* ∈ {1, …, *N*}, the entropy is denoted by :

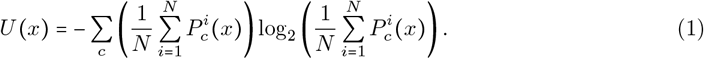

Next, the confidence score at pixel-level was quantified by [28]:

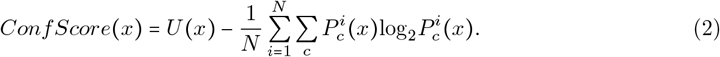

Additionally, a column-wise confidence score for a given column, *col*, was computed where *h* denoted the number of pixels in that column:

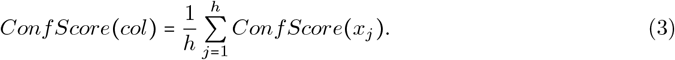

### Prediction process

The prediction process for individual TTE recordings involves several steps, as illustrated in Figure 6 Panel a. M-Mode sections are extracted from the recordings based on the DICOM image metadata. The resulting multi-frame stack is then divided into consecutive, overlapping frames. Each frame was symmetrically zero-padded, centering the original frame within the required 1024 × 1024 pixels canvas. For each frame, the Bayesian U-Net generates the binary segmentation mask *S*_*pred*_ calculated from the averaged predicted probabilities {*P*_*i*_, …, *P*_*N*_ } [28]:

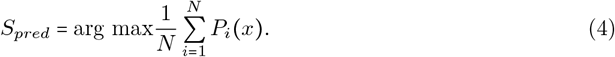

**Figure 6.**
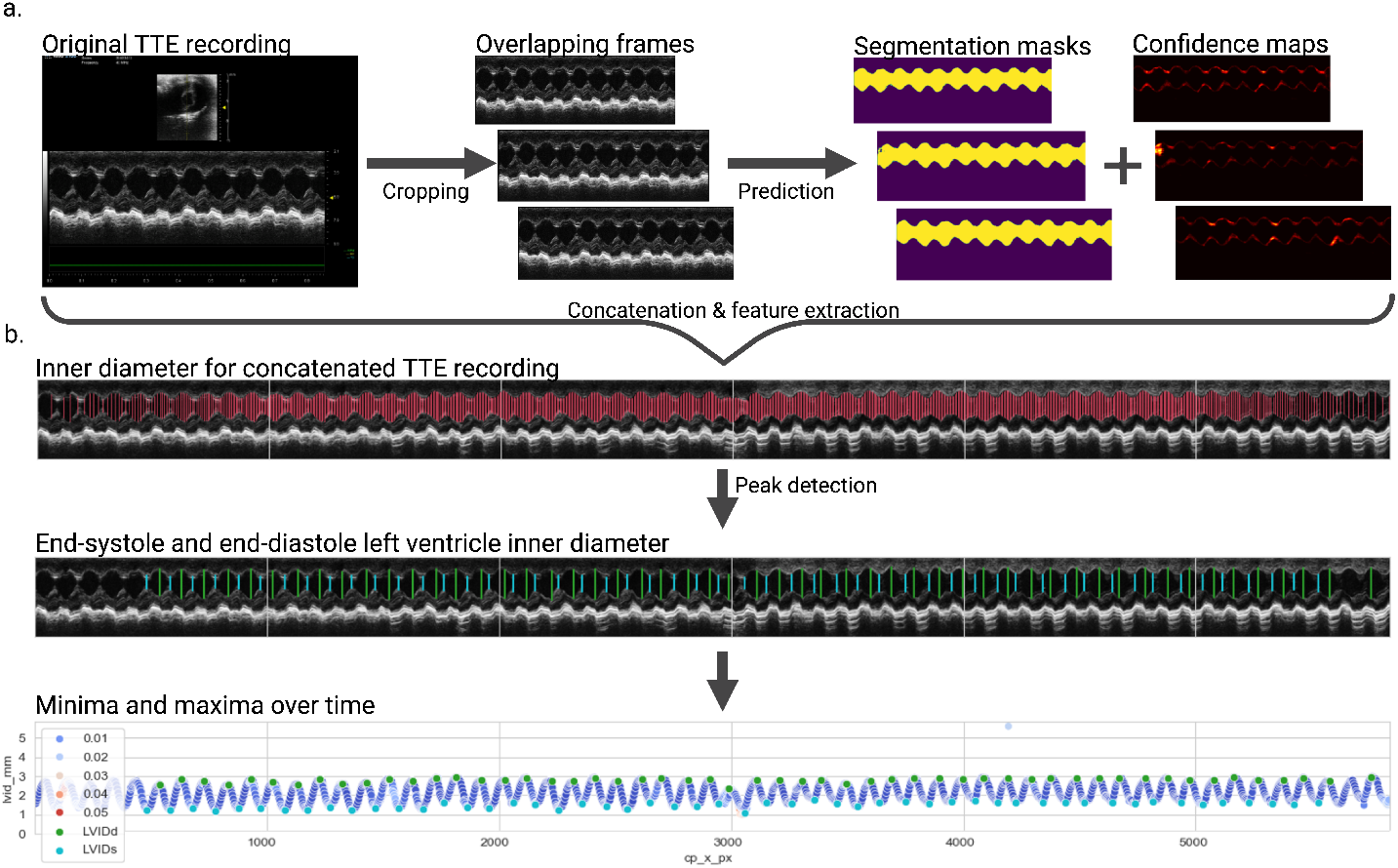
Overview of the data processing workflow in a simplified way. Panel a shows an exemplary TTE recording, which undergoes preprocessing and segmentation prior to feature extraction. After concatenation and temporal alignment, left ventricle inner diameters and their local maxima *LV ID*_*d*_ and minima *LV ID*_*s*_ are derived (panel b). This workflow is followed by the computation of functional diameters, not displayed here.

For uncertainty estimation, the number of Monte Carlo samples *N* was fixed to 10. To assess the model’s confidence throughout the entire frame, the Bayesian U-Net generates confidence maps by aggregating pixel-wise confidence scores *Conf Score x* into an image matching the dimensions of the corresponding frame.

### Extraction of inner diameters

The binary segmentation mask for each frame *S*_*pred*_ was used to determine the inner diameters of the left ventricle in the heart. Beginning from the first pixel column on the left side of each frame, the vertical distance between two pixels, *p*_*l*_ and *p*_*k*_ with *l* < *k*, corresponding to the lower and upper boundaries of the labeled ventricle region, was measured in ascending order. Columns without any pixels classified as part of the left ventricle were skipped. This process was repeated across the entire frame width. To reduce computational load, measurements were taken every fourth pixel column instead of every column.

Next, the diameters measured in pixels were converted to millimeters using the pixel resolutions specified in the DICOM metadata. Each measurement was also assigned a column-wise confidence score *Conf Score (col)* . Two timestamps, *t*_*frame*_ and *t*_*recording*_, were stored for each measurement to indicate its position and ensure temporal alignment across the entire recording.

### Evaluation of the 5-fold cross-validation

To ensure a balanced training dataset and stable model performance, a 5-fold cross-validation was performed. The expert-curated benchmarking dataset was used as test data to evaluate and compare the five resulting models (*M*_*a*_−_*e*_). Once training was completed, each model generated aggregated binary segmentation masks *S*_*pred*_ for the 836 expert-annotated frames based on 10 Monte Carlo samples. The average manually annotated masks *S*_*ann*_ were then compared with the predicted masks *S*_*pred*_ by computing both binary and weighted Dice Scores.

Furthermore, a direct comparison was performed between the inner diameters derived from the two segmentation sources. For this purpose, inner diameters (*LVID*_*ann*_ and *LVID*_*pred*_) were extracted from *S*_*ann*_ and *S*_*pred*_ at identical time points within each frame. The absolute error between annotated and predicted measurements was computed as a mean signed deviation of = *msd LVID*_*pred*_ − *LVID*_*ann*_. The model with the lowest mean signed deviation on the independent test dataset was selected for further predictions.

### Estimation of inner diameters end-systole and end-diastole

To estimate the left ventricle inner diameters at end-systole *LV ID*_*s*_ and end-diastole *LV ID*_*d*_ (Figure 6 Panel b), measurements from concatenated recordings, rather than single frames, were prioritized. These measurements form a time series, maintaining temporal alignment within each recording. In this periodical signal, local minima correspond to the left ventricle inner diameters at end-systole, *min*_*T*_ = *LV ID*_*s*_, whereas local maxima represent the left ventricle inner diameters at end-diastole *max*_*T*_ = *LV ID*_*d*_ at time point *T* .

To avoid misleading outliers - likely caused by incorrect segmentation masks within specific image regions - data points with low confidence scores *Conf Score (col)* were excluded prior to peak detection. Empirical thresholds were derived from outlier analysis stratified by physiological state: *th*_*iso*_ *=* 0.041 for anesthetized mice and *th*_*awk*_ *=* 0.029 for awake mice.

Peak detection was performed using the find_peaks function from scipy v.1.14.1 [60] to identify local maxima and minima in the periodic signal. Parameters were set to *height* = *mean*(*time series*) and *distance =* 125. Local maxima were extracted directly from the time-series data. To identify local minima, the signal was inverted prior to applying peak detection using identical parameter settings.

### Computation of functional parameters

Functional cardiac parameters were derived from left ventricular inner diameters measured at end-systole (*LVID*_*s*_) and end-diastole (*LVID*_*d*_) throughout the entire recording. The temporal variation in heart rate (*HR*) was calculated in beats per minute (bpm) by

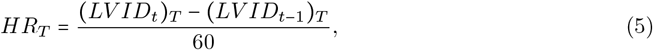

where *T* denotes the time point (in seconds) corresponding to the detected systolic event (local maximum or minimum) measured at the time point *t*. Heart volume at end-systole *Vol*_*s*_ [µl] and end-diastole *Vol*_*d*_ [µl] (6a), ejection fraction *EF* [%] and fractional shortening *FS* [%] (6b), stroke volume *SV* [µl] and cardiac output *CO* [ml/min] (6c) were calculated using the Teichholz formulas [50, 51]:

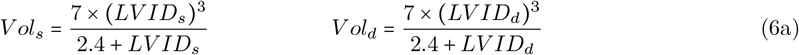

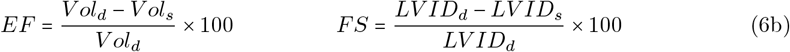

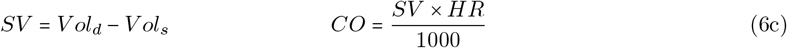

### Data foundation for reanalysis

The dataset used for the reanalysis comprises 65,241 TTE recordings from 18,380 mice across various experimental conditions, including awake and anesthetized mice, as well as mice in early (EA) and late adult (LA) stages, male and female, control and mutant mice (Table 3). This heterogeneity reflects the inherent variability of data acquisition, a common challenge in multi-center studies. The dataset comprised 2±1 recordings per awake mouse and 4±1 per anesthetized mouse. All recordings were systematically screened for duplicate entries. Duplicates were removed once identified.

**Table 3:** Summarized description of the IMPC data used for the reanalysis stratified by anesthetic, sex, and genotype. EA: early adult, LA: late adult.

|  |  |  | EA |  | LA |  |
| --- | --- | --- | --- | --- | --- | --- |
|  |  |  | Recordings | Mice | Recordings | Mice |
| Isoflurane | ♀ | Control | 592 | 304 | 272 | 87 |
|  |  | Mutant | 2222 | 920 | 1240 | 390 |
|  | ♂ | Control | 516 | 292 | 290 | 92 |
|  |  | Mutant | 2082 | 906 | 1117 | 352 |
| Conscious | ♀ | Control | 10355 | 2570 | 448 | 373 |
|  |  | Mutant | 17037 | 4183 | 438 | 359 |
|  | ♂ | Control | 10573 | 2597 | 488 | 395 |
|  |  | Mutant | 17119 | 4178 | 452 | 382 |
| Total |  |  | 60496 | 15950 | 4745 | 2430 |

### Data preparation and QC procedures

All TTE recordings included in the reanalysis were processed using the comprehensive prediction and feature extraction workflow (Figure 6). The results were stored in a MySQL database, and a quality control (QC) procedure was subsequently applied to ensure data integrity and quality compliance.

Manual QC was performed by an IMPC expert for 8,331 TTE recordings from anesthetized mice. During this QC process, *LVID*_*s*_ or *LVID*_*d*_ and their associated functional parameters were excluded if they did not accurately reflect the left ventricular inner dimensions.

Automated quality control instead of manual QC was implemented for 56,910 TTE recordings from awake mice due to the data volume. Here, we targeted high-quality sections in each recording where *LVID*_*s*_ and *LVID*_*d*_ were most densely connected. To this end, the DBSCAN algorithm [33] was applied twice to each recording using the following parameters: *epsilon =* 187.5 and *min*_*samples =* 3. In the first run, DBSCAN identified the most densely packed minima among all *LVID*_*s*_ and categorized them into multiple clusters. Measurements within each cluster were retained, whereas outliers and their corresponding derived cardiac parameters were excluded from subsequent analyses. The procedure was then repeated for *LVID*_*d*_ in the second iteration.

The clustering-based approach effectively filtered out boundary outliers - particularly those at the onset or termination of recordings - that had not been flagged by confidence score thresholding.

### Reference ranges

Reference ranges derived from 23,534 TTE recordings of control mice were established to validate the results of the multivariate clustering approach. These reference ranges were computed for all nine functional parameters and stratified by age, state, and sex. Given that the data did not follow a normal distribution, 95% reference intervals were determined using the 2.5th and 97.5th percentiles, with the median serving as the central estimate. Detailed reference ranges, including medians, percentiles, means, standard deviations, and sample sizes, are provided in the Supplemental Table 2.

### Characterizing abnormal mice

A density-based clustering approach was applied to detect abnormal phenotypes. The dataset was divided into eight subgroups based on age (early and late adult), physiological state (awake and anesthetized), and sex (female and male). Within each subgroup, mutant and control animals were analyzed together. For each mouse, the median and coefficient of variation were calculated across all nine parameters, resulting in an 18-dimensional feature vector. After scaling each subgroup using interquartile ranges, DBSCAN clustering [33] was performed with *min*_*samples =* 19.

Within each subgroup, DBSCAN consistently detected a primary cluster of phenotypically normal animals, whereas mice outside this cluster were treated as outliers and considered abnormal. Mouse cohorts comprising at least four individuals were identified as abnormal if the clustering approach labeled 75% or more of the mutant mice as abnormal. Animals with differing zygosities (homozygous, heterozygous, or hemizygous) but the same single-gene knockout were analyzed separately. The mean cohort size was 4 ± 1, covering 715 genes in total.

Cluster density and sensitivity were fine-tuned by systematically varying the DBSCAN *epsilon* parameter across a pre-defined range for each subgroup (Table 4). Increasing *epsilon* resulted in a sparser cluster of phenotypically normal animals, thereby isolating fewer mice with more pronounced abnormal phenotypes. In contrast, by decreasing *epsilon* the cluster became denser, isolating more animals, including those with milder phenotypic deviations. These observations were further supported by comparisons of functional parameters with their reference ranges, which showed greater deviations at higher *epsilon* values. These systematic differences in cluster density formed the basis for a gene ranking that reflects the consistency and robustness of abnormal assignment across decreasing *epsilon* values.

**Table 4:** Overview of the pre-defined epsilon parameters stratified by age, physiological state, and sex. Setting a lower *epsilon* parameter value leads to a denser major cluster, while setting a higher *epsilon* value results in fewer outlier points and a sparser cluster.

|  |  |  | *Epsilon* |  |  |  |  |
| --- | --- | --- | --- | --- | --- | --- | --- |
|  |  |  | I | II | III | IV | V |
| EA | Isoflurane | ♀<br>♂ | 1.9 | 1.8 | 1.7 | 1.6 | 1.5 |
|  | Conscious | ♀<br>♂ | 1.7 | 1.6 | 1.5 | 1.4 | 1.3 |
| LA | Isoflurane | ♀<br>♂ | 2.0 | 1.9 | 1.8 | 1.7 | 1.6 |
|  | Conscious | ♀<br>♂ | 1.8 | 1.7 | 1.6 | 1.5 | 1.4 |

An additional quality control procedure was conducted to validate the robustness of the gene ranking derived from the clustering analysis. An IMPC expert systematically assessed all 37 ranked genes to evaluate the quality of the derived functional parameters. In total, 22 mice were excluded from analysis because most of their functional parameters were erroneously measured. This exclusion did not affect the data on the gene level. Following this manual QC, 11 genes associated with cardiac phenotypes showed the strongest and most reproducible evidence based on the epsilon value (Table 2). The complete list of all 37 genes linked to cardiac phenotypes is provided in the Supplemental Table 3.

### In-silico analysis of the genes

We compared our TTE findings for genes exhibiting a cardiac phenotype with previously published data. Associations of these genes with cardiac phenotypes were identified through a systematic review and manual annotation of experimental evidence from relevant publications, primarily peer-reviewed small-scale experimental studies. The curated results were integrated into the CIDeR database, a manually maintained multifactorial resource that compiles interactions among heterogeneous factors associated with human diseases [61]. The literature search was completed on 1 November 2025. All supporting references, including PubMed identifiers (PMIDs), are provided in the Supplemental Table 3.

## Data Availabilty

Expert annotations, along with frames extracted from the original TTE recordings, are publicly available at https://doi.org/10.82296/hmgu-nefeli.20exy-nv105. Individual analysis profiles for all mice (*Mybpc3, Acot12, Cep70*) alongside cohort summary plots are provided at https://doi.org/10.82296/hmgu-nefeli.dh2n7-yv037. The original echocardiography dataset supporting this study is available from the corresponding author upon reasonable request.

## Code Availability

We provide a demo application of the analysis workflow, which starts with a TTE recording in DICOM format, continues with frame-wise segmentation, and results in derived cardiac parameters https://github.com/ExperimentalGenetics/EchoVisuALL. The best-performing model is available at https://doi.org/10.82296/hmgu-nefeli.2yphm-15n14.

## Author contributions

Data Analysis: Isabella Galter, Elida Schneltzer; Manuscript Design & Preparation: Isabella Galter; Design & Study Supervision: Elida Schneltzer, Carsten Marr, Nadine Spielmann, Martin Hrabě de Angelis; PI on grant: Martin Hrabě de Angelis and Carsten Marr; Manuscript review: Elida Schneltzer, Nadine Spielmann, Carsten Marr, Martin Hrabě de Angelis.

Members of the IMPC Consortium contributed to data generation and resources.

## Acknowledgments

This work would not have been possible without the support of the IMPC phenotyping centres. Hands-on mouse IMPC phenotyping has been carried out by many laboratory staff with superb experimental skills and unsurpassed dedication. The authors thank these individuals for their contribution to the IMPC data used herein.

- Heaney Jason D. and Christopher S. Ward, Baylor College of Medicine, One Baylor Plaza, Houston, Texas, 77030, United States of America
- Herault Yann and Ghina Bou Abou, Université de Strasbourg, CNRS, INSERM, Institute de la Clinique de la Souris, PHENOMIN, 1 rue Laurent Fries, 67404 ILLKIRCH, France
- Nutter Lauryl M. J., Ann Flenniken and Lois Kelsey, The Hospital for Sick Children, Toronto, ON M5G 1X5, Canada
- Valérie Gailus-Durner, Helmut Fuchs and Ralf Fischer, Institute of Experimental Genetics and German Mouse Clinic, Helmholtz Munich, German Research Center for Environmental Health, Ingolstädter Landstr. 1, 85764 Neuherberg, Germany
- Sedlacek Radislav and Zabrodska Eva, Czech Centre for Phenogenomics, Institute of Molecular Genetics of the Czech Academy of Sciences, Prague, Czech Republic

We gratefully acknowledge the research group, Biocuration for Digital Health (Institute of Experimental Genetics, Helmholtz Munich, German Research Center for Environmental Health, Neuherberg, Germany), led by Andreas Ruepp, including Goar Frischman and Gisela Fobo, for their professional literature research and valuable support of this project.

