## Supplementary figures and images for "EchoVisuALL: From Echocardiography to Gene Discovery"

### Supplemental Figure 1

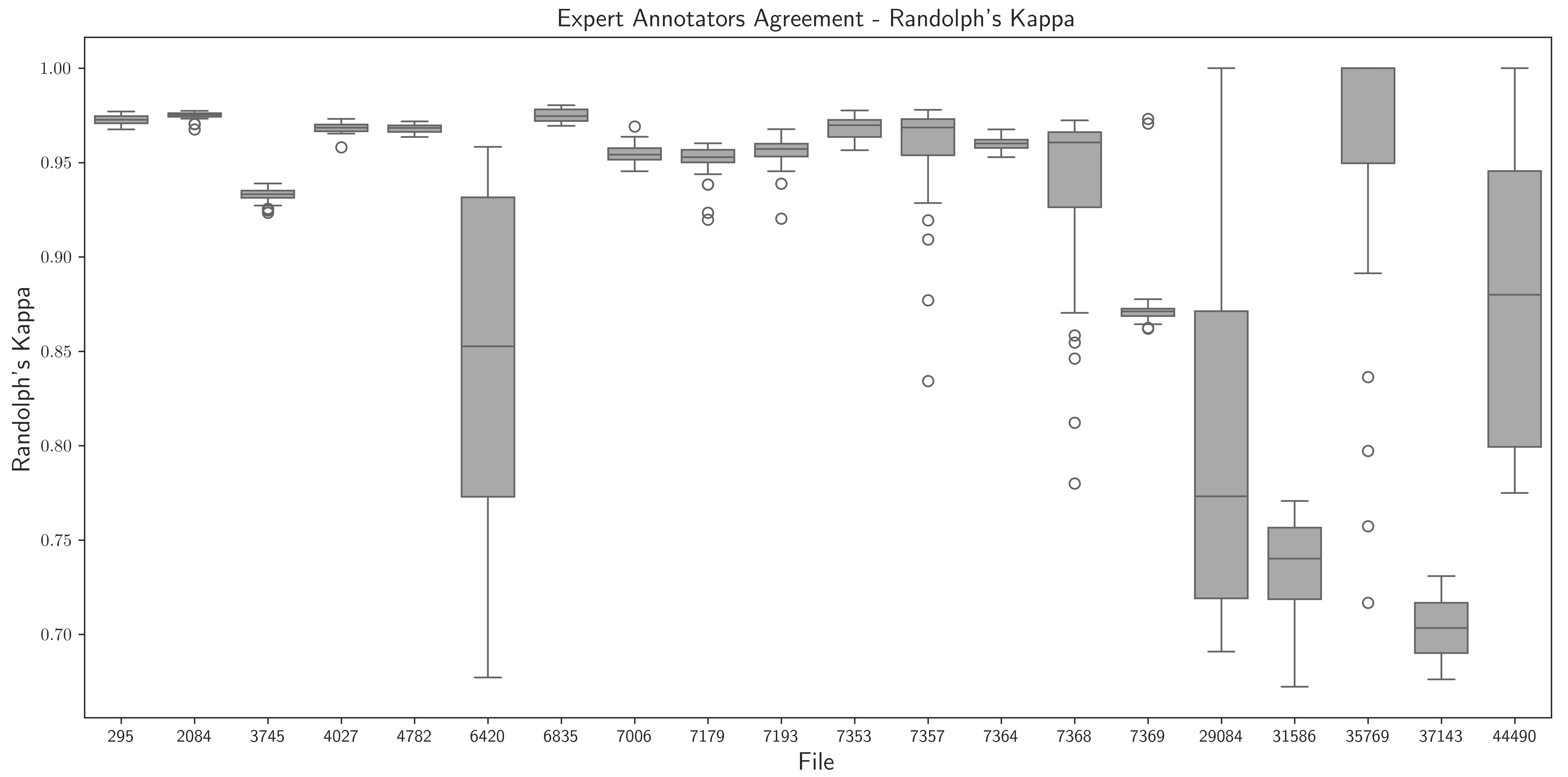
